# An extended DNA-free intranuclear compartment organizes centrosomal microtubules in Plasmodium falciparum

**DOI:** 10.1101/2021.03.12.435157

**Authors:** Caroline S. Simon, Charlotta Funaya, Johanna Bauer, Yannik Voß, Marta Machado, Alexander Penning, Darius Klaschka, Marek Cyrklaff, Juyeop Kim, Markus Ganter, Julien Guizetti

## Abstract

Rapid proliferation of *Plasmodium falciparum* parasites in human red blood cells is the cause of malaria and is underpinned by an unconventional cell division mode, called schizogony. Contrary to model organisms, *P. falciparum* replicates by multiple rounds of closed and asynchronous nuclear divisions that are not interrupted by cytokinesis. Organization and dynamics of the critical nuclear division factors are, however, poorly understood. Centriolar plaques, the centrosomes of *P. falciparum*, are important regulators of division and serve as microtubule organizing centers. Early microscopy studies reveal an acentriolar, amorphous structure although its detailed organization remains elusive. Intranuclear microtubules mediate chromosome segregation, but the small size of parasite nuclei has precluded detailed analysis of their arrangement by classical fluorescence microscopy. We apply recently developed STED, expansion microscopy, and live cell imaging protocols to describe the reconfiguration of microtubules during schizogony. Analysis of centrin, nuclear pore, and microtubule positioning reveals a bipartite organization of the centriolar plaque. While centrin is extranuclear, we confirm by correlative light and electron tomography that microtubules are nucleated in a previously unknown and extended intranuclear compartment, which is devoid of chromatin. This study enables us to build a working model of the organization of an unconventional centrosome and better understand the diversity of eukaryotic cell division modes.

## Introduction

*Plasmodium falciparum* encounters significant population bottlenecks when being transmitted between humans and mosquitoes. To overcome those it undergoes several phases of extensive proliferation. When a mosquito takes up an infected blood meal a rapid series of division events is triggered resulting in the formation of eight male gametes from a single gametocyte within only 15 minutes (Fang et al., 2017; Sinden et al., 1978). During the subsequent oocyst stage in the mosquito midgut thousands of sporozoites are produced of which very few reach the human liver after an infectious mosquito bite (Beier, 1998; Vaughan, 2007). Once those sporozoites invade a hepatocyte they can generate more than ten thousand daughter cells within one cycle, which are then released into the blood (Prudêncio et al., 2006; Sturm et al., 2006). There, repeated rounds of red blood cell invasion, growth, division, and egress cause the high parasite loads, which lead to all clinical symptoms associated with malaria (Schofield, 2007). The cell division processes that underlie these unconventional proliferation events are, however, poorly understood (Francia and Striepen, 2014; Gubbels et al., 2020; Matthews et al., 2018; Simon et al., 2021).

Successful division requires a series of cellular events. Chromosomes must be replicated alongside duplication of the centrosomes, which act as the poles towards which sister chromatids are segregated. Thereafter nuclei are physically separated and the cytoplasm is divided by cytokinesis. *P. falciparum*, however, uses an unconventional division mode, called schizogony where several rounds of nuclear divisions are not interrupted by cytokinesis leading to formation of multinucleated parasite stages (Leete and Rubin, 1996). Although nuclei share a common cytoplasm in schizonts, nuclear divisions are asynchronous (Arnot et al., 2011; Dorin-Semblat et al., 2011; Read et al., 1993). Throughout nuclear division the nuclear envelope remains intact and the DNA is not condensed (Read et al., 1993). Once all rounds of nuclear division are completed each of the 8-28 nuclei are packaged into individual daughter cells, called merozoites (Garg et al., 2015; Reilly et al., 2007; Simon et al., 2021). Upon rupture of the infected host cell, merozoites are released and invade new red blood cells.

Centrosomes are generally regarded as key regulatory hubs of the cell cycle and their duplication limits the number of nuclear divisions (Fu et al., 2015). The centrosome of *P. falciparum* is called centriolar plaque. It exhibits important morphological differences when compared to model organisms such as vertebrate centrosomes or the spindle pole bodies in yeast (Rüthnick and Schiebel, 2018). Available data on the organization of the centriolar plaque is very limited. In early transmission electron microscopy studies, mainly done in oocysts in the mosquito midgut, centriolar plaques appear as electron-dense areas that neither show centrioles nor any other distinct structures (Aikawa et al., 1967; Aikawa and Beaudoin, 1968; Canning and Sinden, 1973; Howells and Davies, 1971; Schrevel et al., 1977; Sinden et al., 1976; Terzakis et al., 1967). Centriolar plaques seem partially embedded in the nuclear membrane, but their positioning relative to the nuclear pore-like “fenestra” remains unclear. Generally, the amorphous appearance of centriolar plaques in electron microscopy has precluded a detailed analysis of their organization so far. The centrosome of a related apicomplexan parasite *Toxoplasma gondii*, which does contain centrioles, shows a bipartite organization with a distinct inner and outer core (Suvorova et al., 2015). Further, few canonical centrosome components are conserved in *Plasmodium* with the exception of centrins, a family of small calcium binding proteins implicated in centrosome duplication (Azimzadeh and Bornens, 2007; Mahajan et al., 2008; Roques et al., 2019), and the microtubule nucleating complex around gamma-tubulin (Zupa et al., 2020). Before we begin to understand the regulation of centriolar plaque duplication and nuclear division we must know the arrangement and dynamics of key division factors around this atypical centrosome.

Centriolar plaques act as microtubule organizing centers. During schizogony intranuclear microtubules organization is very heterogeneous and several atypical structures such as plaques, hemispindles, and the particularly small mitotic spindles have been described (Arnot et al., 2011; Fennell et al., 2006, 2008; Read et al., 1993). Tubulin-rich plaques were equated with centriolar plaques, but their size clearly exceeds the dimensions of electron dense regions described in electron microscopy studies (Gerald et al., 2011). Hemispindles were often interpreted as half-spindles that form a bipolar mitotic spindle by fusion (Fennell et al., 2006, 2008; Read et al., 1993; Schrevel et al., 1977). Inconsistencies in the observed size bring fusion of hemispindles during schizogony into question. While hemispindles observed in electron microscopy are about 0.5 to 0.7 µm in length, which is more consistent with an early stage of a mitotic spindle, hemispindles described by tubulin antibody staining are extensive structures measuring around 2-4 µm. In another study, hemispindles observed in oocysts were interpreted as post-anaphase spindles (Canning and Sinden, 1973). These data provide a controversial view on occurrence, dynamics, and function of hemispindles, which needs to be clarified. We have recently established STED nanoscopy for blood stages, which allowed us to resolve distinct microtubule nucleation sites (Mehnert et al., 2019). These findings were recently confirmed by ultrastructure expansion microscopy (Bertiaux et al., 2021; Guizetti and Frischknecht, 2021). Where microtubule nucleation sites are positioned relative to the nuclear envelope is still an open question.

In this study we use a combination of super-resolution, live cell, and electron microscopy to reveal the ultrastructural organization of the centriolar plaque and microtubules in dividing *P. falciparum* blood stage parasites. We characterize their unconventional dynamics and reveal a novel subnuclear compartment that harbors microtubule nucleation sites and is devoid of chromatin.

## Results

To analyze centriolar plaque and microtubule dynamics we carried out time-lapse imaging of the first nuclear divisions in a blood stage *P. falciparum* parasite strain that episomally expresses PfCentrin1-GFP and was labeled with SPY555-Tubulin, a live cell compatible microtubule dye (Wang et al., 2020). To reduce phototoxicity we applied gentle illumination conditions and used HyVolution-based image processing to generate sufficient image contrast to detect those weak signals (Mov. S1). Initially, microtubules dynamically extended from the centriolar plaque forming hemispindle structures (Fig. 1A). This was followed by a prolonged phase, where the tubulin signal accumulated close to the PfCentrin1-GFP signal. Finally, the tubulin signal elongated while the distance between centrin foci increased, which likely corresponded to anaphase. We quantified the duration of hemispindle, accumulation, and anaphase stages throughout the first three nuclear divisions (Fig. 1B). Every microtubule organization stage was significantly longer in the first division when compared to the second or third (Fig. S1A). The precise time point of initial appearance of the centrin focus varied and was sometimes difficult to determine, which could depend on PfCentrin1-GFP expression levels (Fig. 1C). In some cells appearance of the centrin focus already coincided with the hemispindle microtubule stage (Fig. 1A). In other cells it was only detectable later (Mov. S2) or sometimes not at all. A second centrin focus appeared on average 65 min after the first one (Fig. S1B). This was followed by anaphase indicating that a mitotic spindle was assembled during centrin signal duplication. The now segregated centrin foci were again associated with dynamic hemispindle structures, which subsequently went through a collapsed stage prior to initiating the next duplication and elongation event. Frequently, the second and third round of spindle elongation occurred in an asynchronous fashion.

**Figure 1).**
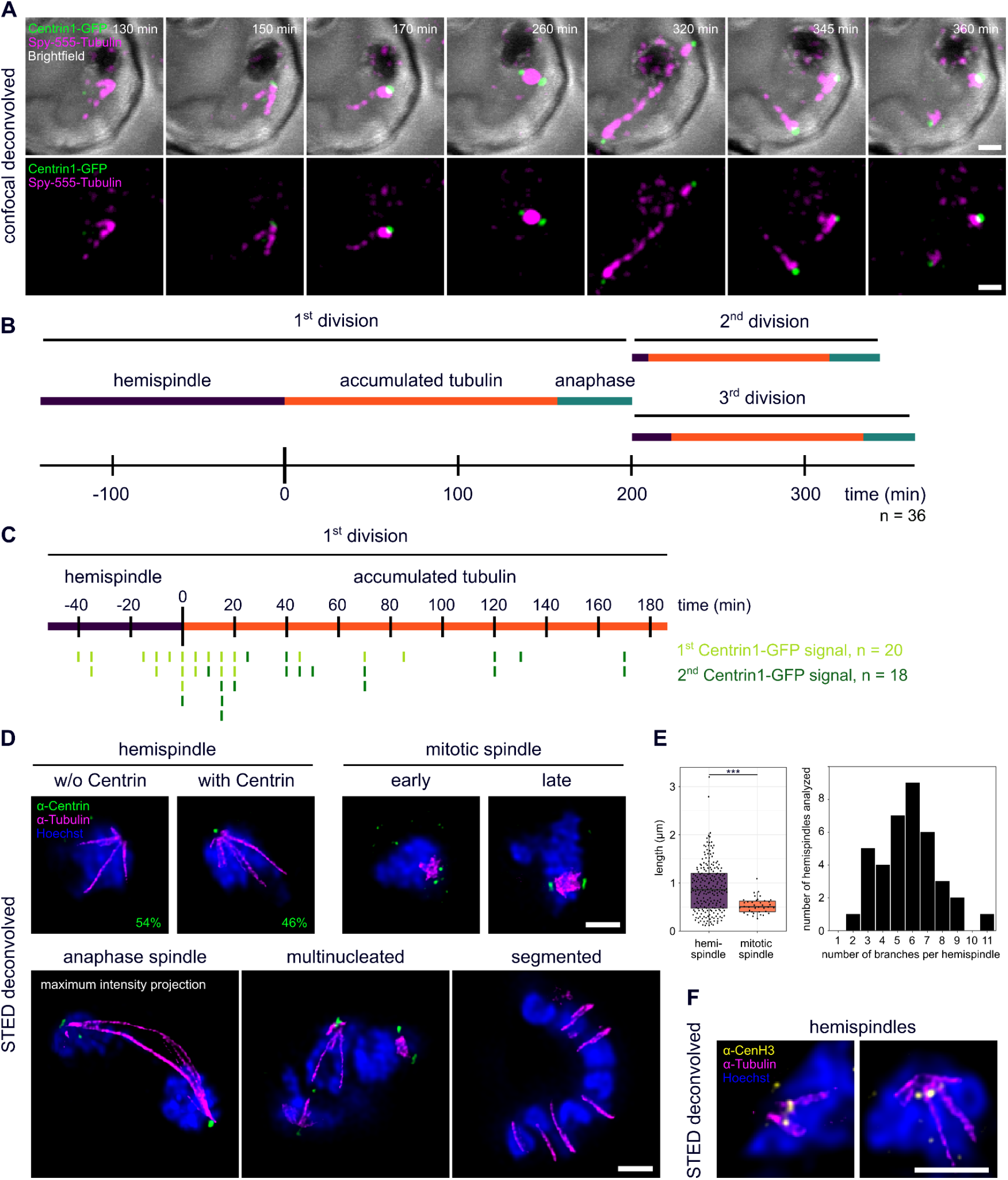
Live cell and super-resolution microscopy of microtubule, centriolar plaque, and centromere reorganization throughout schizogony. A) Deconvolved confocal time-lapse microscopy of first spindle formation and elongation in a schizont ectopically expressing PfCentrin1-GFP (green) and labeled with SPY555-Tubulin (magenta). Maximum intensity projections are shown. B) Quantification of mean duration of three distinct microtubule organization phases in 36 movies. Since most movies (n=32/36) were already started at hemispindle phase we could only quantify the minimal mean length of hemispindle stage for the first division. C) Time points of appearance of first and second clear PfCentrin1-GFP signal normalized to the start of accumulated tubulin phase. D) Dual-color STED nanoscopy images of different schizogony stages of parasites labeled with anti-centrin (green), anti-tubulin (magenta) antibodies and stained with Hoechst (blue). Single slices are shown except for the anaphase spindle. E) Quantification of lengths (905 ±509 nm, n=217) and number of branches per nucleus (5.7 ±2, n=38) and mitotic spindle lengths (535±166 nm, n=38). Statistical difference determined by Welch two-sample t-test. F) like D) with anti-tubulin (magenta) and anti-CenH3 (yellow) showing centromere positioning. All scale bars are 1 µm.

Due to the inherent resolution and sensitivity limits of live cell imaging we could not consistently determine the appearance of the centrin signal and its duplication. Further, we wanted to analyze organization of endogenous centrin and its exact positioning relative to microtubules. Therefore, we imaged parasites at different stages of schizogony after immunolabeling with anti-tubulin and anti-centrin antibodies by dual-color STED nanoscopy (Fig. 1D). While ring and early trophozoite stage parasites do not express tubulin or centrin (Fig. S2), we identified hemispindle structures already in late mononucleated stages. Consistent with the late appearance of PfCentrin1-GFP in our live cell imaging data, only 24 out of 52 analyzed hemispindles in mononucleated cells were associated with an endogenous centrin signal before the first division, while in later stages every nucleus was accompanied by one or two centrin foci. Higher sensitivity of immunofluorescence labeling shows that endogenous centrin accumulates prior to formation of the first mitotic spindle. The collapse of hemispindles into an intense and compact tubulin accumulation was consistently associated with the appearance of two centrin foci. At which point tubulin is reorganized into the bipolar microtubule array, which then forms the mitotic spindle cannot be resolved here. However, early stages where duplicated centrin foci are proximal can be differentiated from late stages where they oppose each other with DNA in the middle. Occasionally, two nuclei connected by an elongated microtubule structure, which we interpret as anaphase spindles, can be observed. In multinucleated stages, several different microtubule organizations could be observed simultaneously in distinct nuclei, reflecting the asynchrony of nuclear divisions. At no point did we detect astral or extranuclear microtubules in schizonts. Cytokinetic segmenter stages on the other hand lack distinct centrin foci and intranuclear microtubules, but clearly display the microtubule cytoskeleton associated with the inner membrane complex.

STED microscopy is, however, partly due to high laser intensities, significantly limited in the acquisition of full z-stacks of entire cells. To reveal the detailed three dimensional organization of spindles in dividing nuclei we therefore employed ultrastructure expansion microscopy (U-ExM), which causes an isotropic expansion of immunolabelled cells (Gambarotto et al., 2019). 3D-rendering of acquired image stacks showed the radial branching of hemispindles (Mov. S3) and the compact organization of mitotic spindles (Mov. S4). These data allowed reliable length measurements of mitotic spindles in 3D averaging at about 560 nm (Fig. 1E). For individual nuclei with hemispindles the length of branches varied substantially between less than 200 nm to more than 3 µm, as did the and number of branches per nucleus, which was ranging from 2 to 11.

Since the role of hemispindles is unclear we wanted to test whether they might be involved in recruitment of centromeres, akin to a “search-and-capture” mechanism (Heald and Khodjakov, 2015), which could assist their clustering at the nuclear periphery (Hoeijmakers et al., 2012). Therefore, we co-labeled tubulin with an anti-CenH3 antibody, which specifically marks centromeric histones (Fig. 1F). As expected, the centromere signal clustered at the periphery next to the centriolar plaque (Zeeshan et al., 2020). However, STED nanoscopy revealed that CenH3 foci were virtually always distinct from hemispindle microtubules, precluding any direct interaction at the acquired time points.

Notably, all images revealed a significant gap between centrin and tubulin signals (Fig. 1D). The question of whether microtubules and centrin locate inside or outside the nucleus remains open. In absence of a known nuclear envelope marker for *Plasmodium* spp., we used a strain where the nuclear pore protein Nup313 had been tagged endogenously with 3xHA (Fig. S3) to partly mark the nuclear boundary (Kehrer et al., 2018). Immunofluorescence co-staining with centrin and tubulin revealed that centrin signals localize on the cytoplasmic side, while microtubule ends are localized inside the nucleus during all stages of schizogony (Fig. 2A). We, also, consistently observed a Hoechst-free region right beneath the centrin foci, which has not been described previously. In nuclei with accumulated tubulin signals those localize within this region. Measurement of this region indicates that its dimensions are not significantly different in hemispindle and mitotic stage nuclei (Fig. 2B).

**Figure 2).**
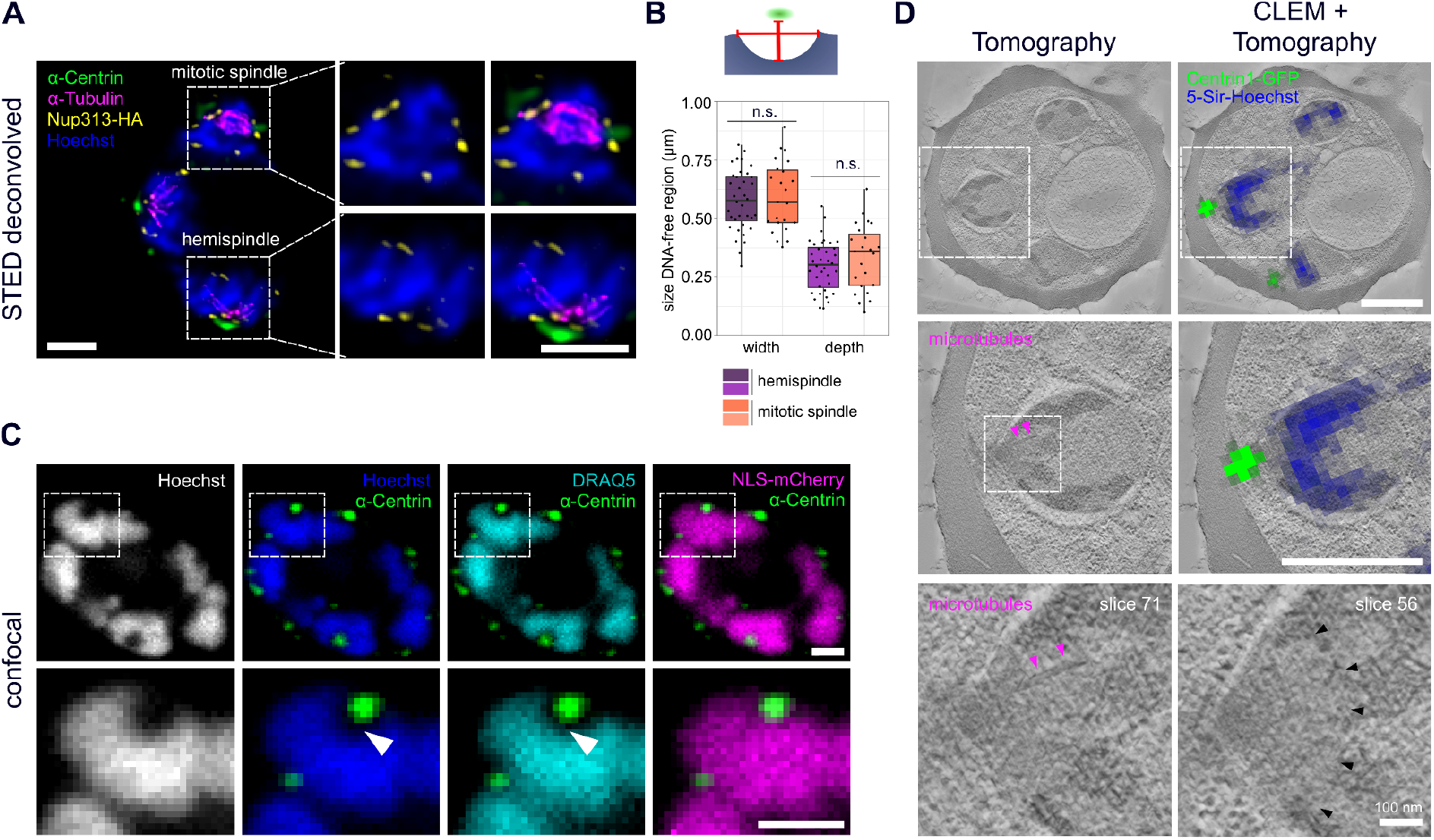
Centriolar plaques are divided in an extranuclear centrin-containing compartment and an intranuclear DNA-free compartment associated with microtubules. A) Dual-color STED nanoscopy images of schizont expressing HA-tagged nuclear pore protein Nup313-HA (yellow) labeled with anti-HA and anti-tubulin (magenta) overlaid with confocal images of anti-centrin (green) and Hoechst-staining (blue). B) Quantification of DNA free region width (hemispindle: 578 ±130 nm, n=36, mitotic spindle: 595 ±148 nm, n=23) and depth (hemispindle: 298 ±108 nm, mitotic spindle: 335 ±141 nm) measured as indicated in schematic. No statistical difference determined by unpaired t-test C) Confocal images of a schizont ectopically expressing NLS-mCherry (magenta) labeled with anti-centrin (green) and stained for DNA with Hoechst (blue) and DRAQ5 (turquoise). D) Correlative in-resin widefield fluorescence and electron tomography images of thick sections of a high-pressure frozen and embedded schizont expressing PfCentrin1-GFP (green) and stained for DNA with 5-SiR-Hoechst (blue). Same cell region containing clear PfCentrin1-GFP foci was imaged by fluorescence microscopy and overlaid with an electron tomogram slice. In zoom-ins, arrows indicate a microtubule (magenta) and a boundary-like region (black) for two tomogram slices. All scale bars are 1µm, except where indicated.

To test if this region is indeed devoid of DNA and not a result of inefficient Hoechst labeling we stained cells with DRAQ5, which is an intercalating DNA dye that is not sensitive to heterochromatin state (Fig. 2C). This staining confirmed the absence of DNA from this region beneath centrin. To further assess the position of the nuclear boundary at the centriolar plaque we used a strain ectopically expressing mCherry tagged with three nuclear localization signals (NLS) to stain the nucleoplasm (Fig. 2C) (Klaus et al., 2021). The NLS-mCherry signal was overlapping with the DNA-free region suggesting that there is an extended subnuclear compartment devoid of DNA associated with the centriolar plaque.

To more directly visualize the nuclear membrane and the ultrastructural features surrounding the centriolar plaque we used electron microscopy (EM). Initial analysis of schizont nuclei with transmission EM suggested that an intranuclear region associated with microtubules is, indeed, delineated by a non-membranous boundary (Fig. S4). However, due to their amorphous structure centriolar plaques cannot always be reliably identified in electron microscopy samples. Hence, we adapted an in-resin correlative light and electron microscopy (CLEM) approach (Kukulski et al., 2012, 2011) to our system using the PfCentrin1-GFP expressing parasite line. In addition, we labeled the cells with the infrared DNA dye 5-SiR-Hoechst (Bucevičius et al., 2019). The fluorescent signal was preserved in the samples prepared for electron microscopy and resin sections were imaged on a widefield fluorescence microscope to identify the residual PfCentrin1-GFP foci (Fig. 2D). Using overview images and finder grids we were able to relocate individual cells at the electron microscope. Overlaying the fluorescence image with the electron tomogram allowed us to unambiguously define the centriolar plaque position. 5-SiR-Hoechst signal was also still detectable after sample preparation but, likely due to imaging a limited section of the nucleus, the staining was not uniform and mostly limited to electron dense heterochromatin regions. The region associated with the centriolar plaque was, although, consistently free from Hoechst staining (Fig. 2D). We could not detect any invagination of the nuclear membrane adjacent to the centrin signal. We, however, identified an underlying region with distinct electron density distribution, which was sometimes associated with microtubules (Fig. 2D). The size and shape of that region corresponded well to the Hoechst-free region observed in our immunofluorescence staining (Fig. 2B) suggesting that this is, indeed, a novel intranuclear compartment with centrosomal function.

Due to difficulties using UV for resin polymerization in the pigmented erythrocytes we also tried embedding in the resin LR gold, which can be polymerized chemically, and these samples were used for the CLEM approach. Still the samples retrieved from UV-polymerized HM20 showed superior contrast and were used for investigating more details of the microtubule organization in nuclei of the schizont stage by electron tomography. We found nuclear stages containing highly elongated individual microtubules which can deform the nuclear envelope at their tips and likely correspond to hemispindles (Fig. 3A & Mov. S5). Here, the sample quality was sufficient to distinctly identify the characteristically shaped microtubule nucleation complex around gamma-tubulin, which demarcates microtubule minus ends (Fig. S5). Those were emerging from discrete positions underlying an electron dense region at the nuclear envelope. A distinct intranuclear compartment could, however, only be surmised in some nuclei when using this sample preparation method (Fig. S6). Mitotic spindles displayed a short but much denser array of microtubules with minus ends clustered at a substantial distance from the nuclear envelope (Fig. 3B & Mov. S6). While this distance in hemispindles ranged from 26 to 96 nm, the distance for the mitotic spindle varied between 88 to 204 nm (Fig. S7). Taken together, these data suggest that intranuclear microtubule nucleation sites are embedded inside an extended amorphous matrix rather than linked to a nuclear membrane-associated centrosomal protein complex.

**Figure 3).**
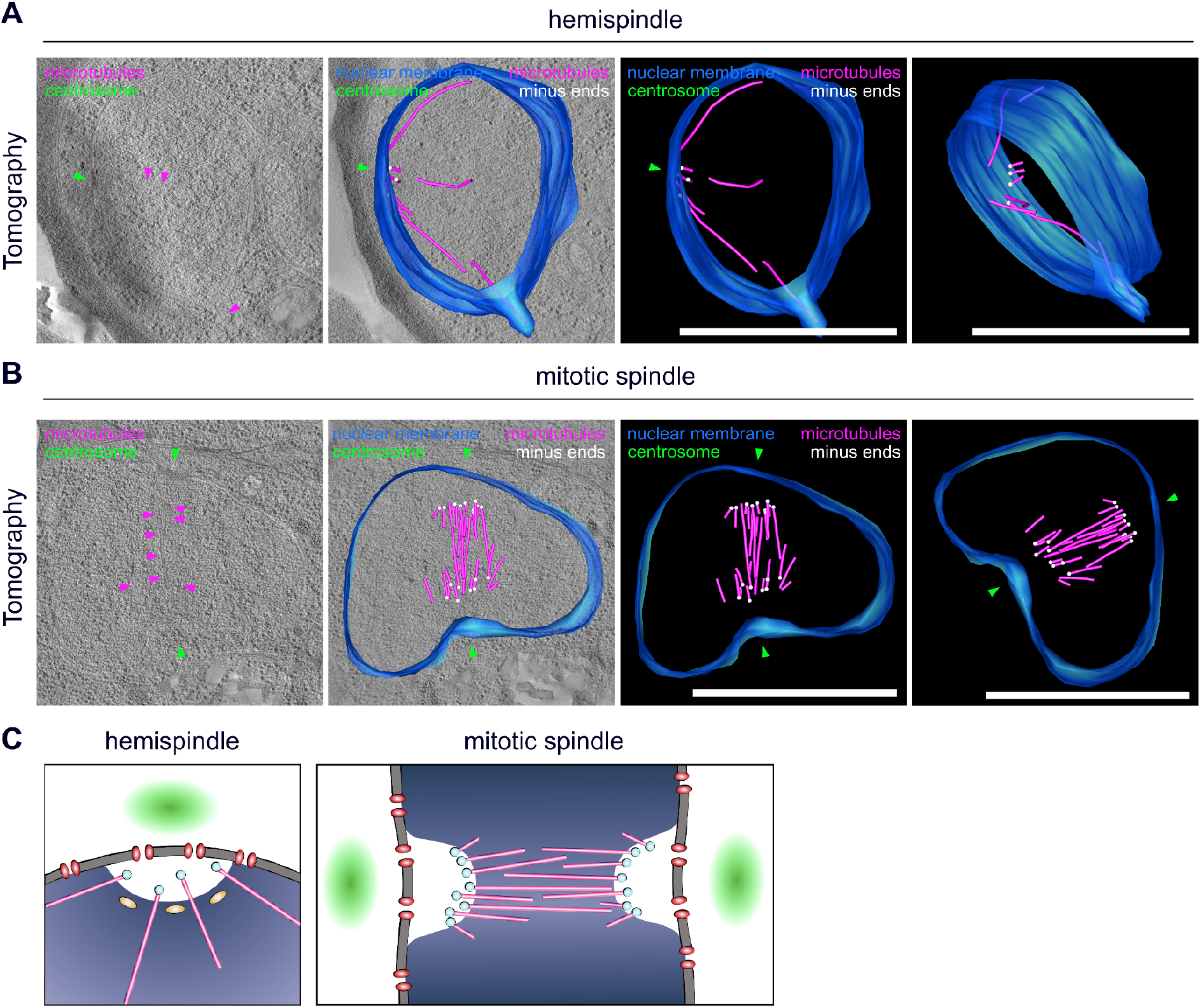
Spindle microtubule nucleation sites are distinct and detached from a defined nuclear membrane associated complex. A) 3D electron tomogram of thick-sections (200nm) of a schizont nucleus in hemispindle stage. Corresponding surface rendering of microtubules (magenta), nuclear membrane (blue), microtubule minus ends (white), and likely position of centriolar plaques (green) are shown. B) as A) for mitotic spindle stage. All scale bars are 1 µm. C) Schematic model of centriolar plaque organization during hemispindle and mitotic spindle stages in blood stage schizonts. Centrin-region (green), nuclear pores (red), microtubules (magenta), microtubule minus ends (pale blue), centromere (yellow), DNA (blue). Content of the DNA-free intranuclear region harboring microtubule nucleation sites is unknown.

## Discussion

Our data provide an entirely novel perspective on the organization of the centriolar plaque and significantly expands on previous concepts depicting it merely as an electron-dense zone, which is inserted in the nuclear membrane (Prensier and Slomianny, 1986). This study highlights the significant differences to the well-characterized spindle pole bodies in budding yeast, where intranuclear microtubules emanate from a small and spatial well-defined protein complex which extends into the cytoplasm (Kilmartin, 2014). We reveal a bipartite organization of the centriolar plaque with an extranuclear region containing centrin and an intranuclear region harboring microtubule nucleation sites (Fig. 3C). While this seems to some degree reminiscent of the inner and outer core described for the centrosome in *T*.*gondii* (Suvorova et al., 2015), we note important differences. In *T. gondii*, centrin has been localized to the inner core and it has been indicated that it is extranuclear (White and Suvorova, 2018). They are likely separated by the prominent additional membrane layer that characterizes the centrocone, but is absent from centriolar plaques. Due to the consistent presence of nuclear pores at the centriolar plaque it is plausible that they are involved in linking the intra- and extranuclear compartments. Whether some of those pores are remodeled to constitute more “fenestra-like” structures remains to be investigated (Bannister et al., 2000).

We further clarify microtubule dynamics, which underlie their heterogeneous organization, previously described in fixed cells (Arnot et al., 2011; Fennell et al., 2008; Read et al., 1993). Hemispindles are already present early in mononucleated cells indicating that they are not per se post-anaphase remnants as suggested earlier (Canning and Sinden, 1973). The lengths and numbers we measured are congruent with recent measurements (Bertiaux et al., 2021), although the slightly higher average number of branches (5.7 vs 4) could be a consequence of us analyzing structures in 3D instead of using projections. Their function, however, remains elusive. We could abate the conspicuous hypothesis that they are directly involved in clustering centromeres at the nuclear periphery, which is consistent with findings in yeast where perinuclear centromere clustering has already been shown to be microtubule-independent (Richmond et al., 2013). An alternative explanation for the presence of hemispindles is that once tubulin accumulates beyond the required critical concentration microtubules polymerize spontaneously (Walker et al., 1988). After the hemispindle stage the tubulin signal collapses into a smaller focus, which is akin to the “tubulin rich plaques” (Gerald et al., 2011). The increased sensitivity and resolution of our assays, however, demonstrate that tubulin accumulations are virtually always associated with two centrin signals indicating that those structures are early mitotic spindles. When exactly microtubules are polymerized into structured bipolar spindles is not resolved, but we can at this point exclude that they are the result of fusion between hemispindles, as previously suggested (Gerald et al., 2011). This early centriolar plaque duplication might be associated with S-phase onset as hypothesized earlier (Francia and Striepen, 2014).

Lastly, we did not observe intranuclear microtubules to be in contact with the nuclear membrane or membrane-associated structures, like for spindle pole bodies (Kilmartin, 2014). Rather, microtubules are nucleated from distinct sites within an extended nuclear compartment which is free of chromatin that has not been described before. In fission yeast a similar configuration has been described although in this case meiotic microtubule arrays emanate from an amorphous compartment just outside the nucleus (Funaya et al,. 2012). The displacement of minus ends away from the nuclear envelope upon mitotic spindle formation, while the dimensions of the centrosomal compartment stay constant, could be attributed to the force generated by the bipolar microtubule array. Despite the absence of centrioles, the preservation of the nuclear membrane, and the lack of conserved factors this amorphous structure is vaguely reminiscent of the pericentriolar material, which makes up the outer layer of vertebrate centrosomes and also harbors microtubule nucleation complexes (Woodruff et al., 2014). Deciphering the composition of this novel nuclear compartment, understanding how it is assembled, and how it splits are some of the more pressing questions emerging from this study.

## Materials and Methods

### Parasite culture

*Plasmodium falciparum* cell lines NF54_Centrin1-GFP and 3D7_Nup313-HA_glms and NF54 wild type were cultured in O+ human red blood cells in RPMI 1640 medium supplemented with 0.2 mM hypoxanthine, 25mM HEPES, 0.5% Albumax and 12.5 µg/ml Gentamicin. Cultures were maintained at a hematocrit of about 3% and a parasitemia of 3-5%. Parasite cultures were incubated at 37°C with 90% humidity, 5% O_2_ and 3% CO_2_. To synchronize cultures, late stage parasites were lysed by 5% sorbitol treatment.

### Plasmid constructs

To generate a pArl-Centrin1-GFP plasmid for episomal expression of PfCentrin1-GFP in NF54, we used a pArl-PfCentrin3-GFP plasmid (kindly provided by Tim Gilberger). The pArl-PfCentrin3-GFP plasmid was digested with KpnI-HF and AvRII to cut out the PfCentrin3. The PfCentrin1 insert sequence was generated from *P. falciparum* cDNA via PCR using the forward primer CGACCCGGGATGGTACCATGAGCAGAAAAAATCAAACTATG and reverse primer TTCTTCTCCTTTACTCCTAGGAAATAAGTTGGTCTTTTTCATAATTC. PfCentrin1 and the digested backbone were ligated using Gibson Assembly and the sequence was verified by Sanger sequencing. To generate the pSLI-Nup313-3xHA_glms construct, first the HA and glmS sequences were ligated into the pSLI-TGD plasmid, a kind gift of Tobias Spielmann (Birnbaum et al., 2017), to obtain the pSLI-3xHA-glmS. The glmS ribozyme was amplified from the plasmid pARL_glmS (a kind gift from Jude Przyborski) and inserted by Gibson assembly into pSLI-TGD, downstream of the NeoR/KanR resistant cassette, using primers 0079 and 0082. For the 3xHA tagging, its sequence was first PCR-amplified with primers 0126 and 0127 from pDC2-cam-coCas9-U6.2-hDHFR (a kind gift from Markus Lee) and cloned by Gibson assembly into the NotI and MluI digested pSLI-glmS plasmid. Lastly, 757bp of NUP313 genomic sequence (without stop codon) was PCR amplified using primers 0163for and 0164rev and cloned into the modified pSLI-3xHA-glmS plasmid using NotI and Mlu1 restriction sites. The p3-NLS-L3-mCherry construct was obtained from the p3-NLS-FRB-mCherry plasmid, a kind gift of Tobias Spielmann (Birnbaum et al., 2017), by removing the FRB domain with NheI and KpnI digestion and consecutive ligation. Correct sequence of inserts was verified by Sanger sequencing

### Parasite transfection

Transgenic parasites were generated by electroporation of sorbitol-synchronized ring-stage parasites with 50-100 μg of purified plasmid DNA (Qiagen). To select for the plasmids pArl-PfCentrin1-GFP and p3-NLS-L3-mCherry, we used 2.5 nM WR99210 (Jacobus Pharmaceuticals) or 5 μg/ml blasticidin S (InVivoGen), respectively. To select for integration of the Nup313-HA_glms construct into the genome, we followed the protocol published previously (Birnbaum et al., 2017), using 800ug/ml Geneticin-G418 (Thermo Fisher Scientific). PCRs across the integration junctions and testing for leftover unmodified locus to exclude the presence of wild type were performed (Fig. S3). Limiting dilution was done to obtain clonal parasite lines.

### Seeding of infected red blood cells on imaging dishes

For live cell imaging, cells were seeded on round imaging dishes with glass bottom (µ-Dish 35 mm, ibidi), for immunofluorescence stainings on 8-well chambered glass slides (µ-Slide 8 Well, ibidi) as described previously (Gruring et al., 2011; Mehnert et al., 2019). Briefly, the glass surface was coated with Concanavalin A (Sigma, 5 mg/ml in water) for 20-30min at 37°C. Dishes were rinsed twice with prewarmed incomplete RPMI 1640 medium lacking Albumax and hypoxanthine. Infected erythrocyte culture (500 µl for 35mm dishes, 150µl for each well of 8-well glass slides) was washed twice with incomplete medium by centrifugation (1000g, 30sec), before addition of the cells onto the glass. Cells were allowed to settle for 10min at 37°C. By gentle shaking and washing of the glass slides with prewarmed incomplete medium, unbound cells were removed until a monolayer of red blood cells remained on the glass surface. Incomplete medium was replaced by complete medium (4 ml for 35mm dishes, 200 µl for each well of 8-well glass slides) and cells were maintained in the incubator until they were prepared for live cell imaging or fixed for immunofluorescence staining.

### Immunofluorescence assay

Immunofluorescence staining for confocal and STED microscopy was performed as described previously (Mehnert et al., 2019). Briefly, after seeding cells at ring stage, parasites were fixed in schizont stages with prewarmed 4% PFA/PBS for 20min at 37°C. PFA was washed off twice with PBS. Fixed cells were either stored in PBS at 4°C for later immunofluorescence staining, or stained immediately. First, cells were permeabilized with 0.1% Triton X-100/PBS for 15min at room temperature and rinsed three times with PBS. To quench free aldehyde groups, cells were incubated with freshly prepared 0.1 mg/ml NaBH4/PBS solution for 10min. Cells were rinsed thrice with PBS and blocked with 3% BSA/PBS for 30min. In the meantime, primary antibodies were diluted in 3% BSA/PBS and centrifuged at 14.800 rpm for 10 min at 4°C to remove potential aggregates. Cells were incubated with primary antibodies (see below) for 2h at room temperature. Next, cells were washed three times with 0.5% Tween-20/PBS. Incubation with secondary antibodies (see below) plus Hoechst in 3% BSA/PBS was performed for 1h preceding removal of aggregates as described for primary antibodies. After washing twice with 0.5% Tween-20/PBS and once with PBS, cells were stored in PBS at 4°C in the dark until imaging. For longer storage, antibodies were occasionally fixed after staining with 4% PFA/PBS for 10-15min at room temperature. After washing thrice with PBS, cells were likewise stored in PBS at 4°C in the dark.

### Antibodies

To stain microtubules, the mouse anti-alpha-tubulin B-5-1-2 was used at a dilution of 1:500 (Sigma, T5168). Polyclonal rabbit anti-TgCentrin1 (Fig. 2B) antibody was a kind gift of Marc-Jan Gubbels and applied at a 1:2000 dilution. Polyclonal rabbit anti-CenH3 antibody was a kind gift of Alan Cowman and diluted 1:200 (Volz et al., 2010). Rat anti-HA 3F10 (Sigma) was applied at a 1:500 dilution. To generate a polyclonal rabbit anti-PfCentrin3 antibody a codon-optimized sequence of PfCentrin3 (PF3D7_1027700.1) was synthesized (ThermoFisher, GeneArt Strings), and cloned into the pZE13d vector (Lutz and Bujard, 1997) with an N-terminal 6xHis tag via Gibson assembly (Hifi DNA Assembly, NEB) using ClaI and PstI restriction sites. The construct was transformed into chemically competent W3110Z1 E.coli and colonies were inoculated in 800 ml LB-Amp expression culture, which was incubated at 37°C and 130 rpm until reaching an OD600 of 0.5. After induction with 1 mM IPTG, incubation continued for 3 h after which harvested bacteria were lysed via sonication. The lysate was cleared via centrifugation and recombinant PfCen3-6xHis were purified from the soluble fraction using Ni-NTA agarose beads (QiaGen) according to the manufacturer’s recommendations. The buffer was exchanged to PBS via overnight dialysis and the protein further purified using the Superdex 75 10/300 size exclusion column (Cytiva). The final protein was used for a 63-day rabbit immunization regimen and affinity purification of the resulting serum performed by Davids-Biotechnology, Regensburg. The secondary antibodies used for STED, namely anti-mouse-STAR580 (Abberior), anti-rabbit-Atto647 (Sigma), anti-rabbit-Atto594 (Sigma), anti-mouse-Atto647 (Sigma) and anti-rat-Atto594 (Biomol) were all used at a 1:200 dilution. The secondary antibodies anti-rabbit Alexa Fluor Plus 488 (Thermo Fisher Scientific) and anti-rabbit Alexa488 (Thermo Fisher Scientific) were diluted 1:1000. To stain DNA, DRAQ5 (Biostatus) or Hoechst33342 (Thermo Fisher Scientific) were used (both 1:1000). To increase the signal of 3xNLS-mCherry (Fig. 2B), cells were incubated with RFP-Booster nanobody coupled to Atto 594 (Chromotek) at a dilution of 1:200.

### Preparation of infected red blood cells for live cell imaging

For live cell imaging, NF54_PfCentrin1-GFP cells were seeded on glass bottom dishes as described above. Imaging medium, i.e. phenol red-free RPMI 1640 supplemented with stable Glutamine and 2 g/l NaHCO3 (PAN Biotech) with all other supplements as in the parasite culture medium, was equilibrated in the cell culture incubator for several hours. Immediately before imaging, 9 ml of equilibrated imaging medium were supplemented with 4.5 µl (1:2000 dilution) of the live microtubule dye SPY555-Tubulin (Spirochrome). Culture medium in the glass bottom dish with seeded cells was replaced by 8 ml imaging medium, closed tightly without creating air bubbles, and sealed completely with parafilm. The imaging dish was directly taken to the incubation chamber of the microscope, prewarmed to 37°C.

### Super-resolution confocal and STED microscopy

Confocal microscopy of fixed and live cells was performed on a Leica TCS SP8 scanning confocal microscope with Lightning (LNG) module. LNG enables automated adaptive deconvolution after acquisition to generate super-resolution images. All images were acquired using a 63x 1.4 NA objective, GaAsP hybrid detectors and spectral emission filters. For live cell imaging, the adaptive lightning acquisition mode was used with a pinhole of 1.2 airy units resulting in a pixel size of 53.8 nm and a total image size of 18.45 × 18.45 µm (344 × 344 pixels). The pixel dwell time was 488ns. Every 5min, a z-stack was taken of each cell with a total size of 6 µm and an z-interval of 0.5 µm. Centrin1-GFP was excited with a 488nm laser at a laser power of 0.5%, SPY555-Tubulin was excited with a 561nm laser at a laser power of 2%. Cells were imaged over night for a maximum of 13h. For confocal imaging of fixed cells, the LNG mode was turned off and cells were acquired using a pinhole of 1 airy unit, a pixel size of 72.6 nm and a total image size of 9.3 × 9.3 µm (128 × 128 pixels). The pixel dwell time was 488ns. Z-stacks of 6.27 µm were acquired with an z-interval of 0.3 µm. RescueSTED microscopy was performed on a single-point scanning STED/RESOLFT super resolution microscope (Abberior Instruments GmbH), equipped with a pulsed 775 nm STED depletion laser and three avalanche photodiodes for detection. Super-resolution images were acquired with a 100x 1.4NA objective, a pixel size of 20 nm and a pixel dwell time of 10 µs. The STED laser power was set to 10-20%, while the other lasers (488, 594 and 640) were adjusted to the antibody combinations used. To prevent destruction of hemozoin-containing cells by the high-intensity STED laser, intensity thresholds (CONF levels) were defined, which needed to be reached in a confocal image before automatic activation of the STED laser (adaptive illumination). CONF levels varied between 10 and 110 and were adjusted individually for every cell. To acquire z-stacks (anaphase spindle, figure 1B), a total z-stack of 3.9 µm was acquired using a z-step size of 300nm.

### Ultrastructure Expansion Microscopy (U-ExM)

U-ExM was performed as described previously (Gambarotto et al., 2021, 2019), with slight modifications. Schizont parasite pellet was enriched using QuadroMACS Separator (Miltenyi) and added to a Poly-D-Lysin-coated coverslip to settle for 10 min at 37 °C. Excess liquid was removed, and cells were fixed with prewarmed 4 % PFA/PBS for 20 min at 37 °C. Cells were washed three times with prewarmed PBS, and the coverslip was transferred into a 6-well plate. After removal of PBS, the well was filled with 1 ml 1.4 % Formaldehyde (Sigma) / 2% Acrylamide (Sigma) in PBS and incubated for 5 h at 37 °C. Sodium acrylate (Santa Cruz Biotechnology, 7446-81-3) was solubilized in Milli-Q water. Protein denaturation was prolonged to 90 min at 95 °C. The first round of expansion was performed in Milli-Q water for 30 min, before water was changed for overnight incubation. Gel was washed 2x 15 min with PBS followed by blocking for 30 min with 3% BSA/PBS. As primary antibodies, mouse anti-alpha-tubulin B-5-1-2 (Sigma, T5168), mouse anti-alpha-tubulin TAT-1 (Sigma, 00020911), mouse anti-beta-tubulin KMX-1 (Sigma, MAB3408), rabbit anti-PfCentrin3 and Rat anti-HA 3F10 (Sigma) were diluted 1:250 in 1.2 ml 3% BSA/PBS. Solution was spun down for 10 min with 21.1 g at 4 °C to remove aggregates. Gel was incubated with antibodies for 2 h 45 min at 37 °C with agitation. The gel was washed 5 × 10 min with 2 ml 0.5% Tween-20/PBS. As secondary antibodies, anti-mouse-STAR580 (Abberior) and anti-rabbit-Atto647 (Sigma) were diluted 1:100, and anti-Rat-Alexa488 (Thermo) was diluted 1:500 in 1.2 ml 3% BSA/PBS. Hoechst33342 (Thermo) was added at 1:100 dilution, and the solution was spun down as previously described. Incubation was performed at 37 °C for 2 h 30 min with agitation. The gel was washed 5 × 10 min with 2 ml 0.5% Tween-20/PBS afterwards. The second round of expansion was performed as described. Samples were imaged on Leica SP8 in standard confocal mode as described above, with a pixel size of 72.22 nm. Image analysis was performed as described above, and 3D movies were rendered using Imaris (Oxford Instruments).

### Image analysis and quantification

Most images were analyzed using Fiji (Schindelin et al., 2012). Quantification of time-lapse images was performed on images after LNG adaptive deconvolution. Therefore, cells were examined manually to determine the changes of individual tubulin stages over time as well as the first stable appearance of the centrin signal. All deconvolved images shown were deconvolved using Huygens professional using express deconvolution with the standard template. Quantification of hemispindle and mitotic spindle length in U-ExM samples were measured using 3D-distance measurement tools in Imaris and corrected by the expansion factor of 4.5x. Dimensions of DNA-free regions were measured in cells acquired with LNG-mode on the Leica SP8 slices, where the region underlying a centrin signal was visible from the side. 3D distances between MT ends and nuclear membrane were measured in the segmented tomography model using the mtk program in the IMOD software package. Data analysis and depiction were performed using Excel and R studio.

### Preparation of infected RBCs for electron tomography

For high pressure freezing of infected erythrocytes, late-stage parasites of the NF54_Centrin1-GFP strain (2-4 ml packed erythrocytes in culture, 3-5% parasitemia) were purified using magnetic activated cell sorting (VarioMACS™ Separator, Miltenyi Biotec). Importantly, schizonts were not in contact with PBS before HPF, as we have shown recently that hemispindle microtubules are not detectable when parasites were fixed immediately after PBS incubation (Mehnert et al., 2019). For high pressure freezing, around 1.5 µl of concentrated purified schizont pellet was transferred into aluminium or gold carriers (3 mm diameter, 100 or 200 µm depth; Leica Microsystems) and high pressure frozen with EM ice (Leica Microsystems). Freeze substitution was done in a Leica EM AFS2 (Leica Microsystems). Samples were freeze substituted in 0.3% uranyl acetate in dry acetone for 24h at -90°C, followed by an increase in temperature from -90°C to -45°C in 9h (5°C/h). Samples were incubated for another 5h at -45°C, before rinsing 3 × 10 min with dry acetone. Acetone was replaced by increasing concentrations of the Lowicryl HM20 (25%, 50% and 75%) in dry acetone at -45°C for 2h each. Cells were incubated in 100% HM20 at -45°C, after 12h the solution was again replaced by 100% HM20 and incubated for another 2h at

-45°C. To polymerize HM20 and therefore embed the samples in the resin, UV light was applied for 48h at -45°C, for another 13h while increasing the temperature from -45°C to +20°C (5°C/h) and 48h at 20°C. The RBC pellets were not well polymerized due to the pigmentation of the cells and areas with well-embedded cells had to be selected for sectioning. Polymerized cells were trimmed and sectioned on a UC7 ultramicrotome (Leica Microsystems). 200 nm-thick sections were collected on Formvar-coated copper slot grids and contrasted with 3% uranyl acetate and Reynold‘s lead citrate. Sample quality was checked on a Jeol JEM-1400 80 kV transmission electron microscope equipped with a 4k by 4k pixel TemCam F416 digital camera (TVIPS). For image acquisition, the EM-Menu (TVIPS) software was used. For tomography, sections were placed in a high-tilt holder (Model 2040; Fischione Instruments; Corporate Circle, PA) and the cells were recorded on a Tecnai F20 EM (FEI, Eindhoven, The Netherlands) operating at 200kV using the SerialEM software package (Mastronarde, 2005). Images were taken every degree over a ±60° range on an FEI Eagle 4K × 4K CCD camera at a magnification of 19000x and a binning of 2 (pixel size 1.13 nm). The tilted images were aligned using tilt series patch tracking. The tomograms were generated using the R-weighted back-projection algorithm. To reconstruct the complete hemispindle, tomograms were collected from 3 serial sections, aligned and joined by using the eTomo graphical user interphase (Höög et al., 2007). Tomograms were displayed as slices one voxel thick, modelled, and analyzed with the IMOD software package (Kremer et al., 1996). Capped ends of microtubules were identified as minus ends in accordance to earlier microtubule studies with similar preservation techniques (Gibeaux et al., 2012; Höög et al., 2007; O’Toole et al., 2003).

### Correlative in-resin widefield fluorescence and electron tomography (CLEM)

For in-resin CLEM, magnetically purified NF54_Centrin1-GFP late stage parasites were incubated with 1 µM of the live dye 5-SiR-Hoechst for about 1h at 37°C. HPF was performed as described above. Freeze substitution and embedding were done in an Automatic Freeze Substitution System (AFS2, Leica Microsystems), but pipetting steps were performed manually. Cells were freeze-substituted in 0.3% uranyl acetate in dry acetone for 29h at -90°C, before the temperature was increased to -45°C in 9h (5°C/h) and kept at -45°C for at least 5h. The freeze substitution solution was replaced by 100% cold, dry ethanol and the temperature was increased from -45°C to -25°C in 1h (20°C/h). Samples were incubated with increasing concentrations (25%, 50% and 75%) of LRGold (London Resin company) in dry ethanol at -25°C for 2h each. Afterwards, 100% LRGold was added, removed, and again added for overnight incubation of the samples at -25°C. The resin LRGold was supplemented with the initiator, 1.5% benzoyl peroxide, on ice. The solution was inverted carefully to prevent oxygen incorporation and immediately placed at -20° to prevent direct polymerization. Samples were incubated with 100% LRGold with initiator for 26h at -25°C. Temperature was increased from -25°C to 20°C in 9h (5°C/h) and samples stayed at 20°C for 24h for full polymerization. 300nm-thick sections were cut using a UC7 ultramicrotome (Leica Microsystems) and collected on Formvar-coated finder grids. Immediately after sectioning, the grid was placed in a drop of PBS with pH 8.4 (Ader and Kukulski, 2017) on a glass coverslip with the sections facing the bottom. A second glass coverslip was added on top and the sandwich was mounted in a metal ring holder (Kukulski et al., 2011). Fluorescence was imaged directly on a Zeiss Axio Observer.Z1 widefield system equipped with an AxioCam MR R3 camera and a 63x oil objective (1.4 numerical aperture). For excitation of Centrin1-GFP, a 488nm laser was used, for excitation of 5-Sir-Hoechst, a 587nm laser, both set to a laser power of 95% and an exposure time of 900ms. Images were taken with a pixel size of 102nm and a total size of 1388×1040 pixels per tile scan. Subsequently, the sections were contrasted with 3% uranyl acetate and Reynold’s lead citrate. After checking the ultrastructure quality of the Centrin1-GFP positive cells on a Jeol JEM-1400 80 kV transmission electron microscope (Jeol), tomography was performed on a Tecnai F30 EM TEM (EMBL Heidelberg), operating at 300kV. Tilt series of 200 nm thick sections were recorded at the range of +-60° with 2° interval, on a 4x4k CCD camera (Gatan - OneView), using Serial-EM acquisition software. 3D reconstructions and further analyses were conducted using “etomo” Image Processing Package (Boulders, Colorado). Correlation of fluorescence and electron tomography images was performed manually using Fiji, GIMP 2.10.20. and Inkscape.

### Transmission electron microscopy of Spurr-embedded infected RBCs

For high pressure freezing, erythrocytes infected with NF54 wild type parasites (500µl packed red blood cells in culture, 6% parasitemia) were purified using magnetic activated cell sorting (QuadroMACS™, Miltenyi Biotec). Purified late stages were accompanied with 30 µl uninfected red blood cells and cultured for 4h to let the cells recover. High-pressure freezing using Leica EM ICE (Leica Microsystems) was performed as described above. Freeze substitution was done in an Automatic Freeze Substitution System (AFS2, Leica Microsystems). Samples were freeze substituted in 0.2% Osmium tetroxide, 0.3% uranyl acetate and 5% H_2_O in dry acetone for 1h at -90°C. Temperature was increased from -90°C to +20°C in 22h (5°C/h) and samples stayed at 20°C until further processing. Next, samples were washed three times with dry acetone. The pellets detached from the carriers and were combined in a 1.5 ml reaction tube in dry acetone and pelleted for 2min, 2200rpm. Acetone was replaced by a 1:1 mixture of Spurr’s resin (Serva, Heidelberg) and dry acetone. After 2h incubation at room temperature, the mixture was replaced by 100% Spurr’s resin and incubated over night at room temperature. The Spurrr’s resin was removed and again replaced by 100% Spurr’s resin. Polymerization of the samples was performed at 60°C for 1-2 days. Samples were trimmed with a UC7 ultramicrotome (Leica Microsystems) and 70nm thin sections collected on Formvar-coated slot grids. Images were taken on a Jeol JEM-1400 80 kV transmission electron microscope (Jeol) equipped with a 4k by 4k pixel TemCam F416 digital camera (TVIPS). For image acquisition, the EM-Menu (TVIPS) software was used.

## Author Contributions

Experiments were planned and designed by C.S.S with the help of J.G. Light microscopy and immunofluorescence assays were carried out and analyzed by C.S.S. Electron microscopy was realized by C.S.S with help by C.F. CLEM was realized by C.S.S with help by C.F, M.C and J.K. Expansion microscopy was carried out by C.S.S and J.B. PfCentrin1-GFP strains and the anti-PfCentrin3 antibody were produced by Y.V. NLS-mCherry and Nup313-HA strains were generated by D.K, M.M and A.P with the help of

M.G. J.G and C.S.S wrote the manuscript with comments by the other authors.

## Acknowledgments

We thank: The Infectious Diseases Imaging Platform for imaging support (idip-heidelberg.org). The Electron Microscopy Core Facility (EMCF) at Heidelberg University for providing electron microscopy services. PlasmoDB for their *Plasmodium* Informatics Resources. Stefan Pitsch and Luc Reymond (Spirochrome LTD) for providing their SPY555-Tubulin dye. Grazvydas Lukinavicius and Jonas Bucevicius (Max Planck Institute for Biophysical Chemistry) for providing the 5-SiR-Hoechst dye. Paul Guichard (University of Geneva) for sharing their U-ExM protocol. Nicolas Lichti for help with molecular cloning. Marc-Jan Gubbels (Boston College) for the anti-TgCentrin1 and Alan Cowman (WEHI Melbourne) for the anti-CenH3 antibodies.

