## Supplementary figures and movie legends for "An extended DNA-free intranuclear compartment organizes centrosomal microtubules in Plasmodium falciparum"

### Supplementary Data

**Supplementary movies** are deposited under:
https://heibox.uni-heidelberg.de/d/dcbfdb88b6e7492fbfaa/
Corresponding legends can be found at the bottom of this document.

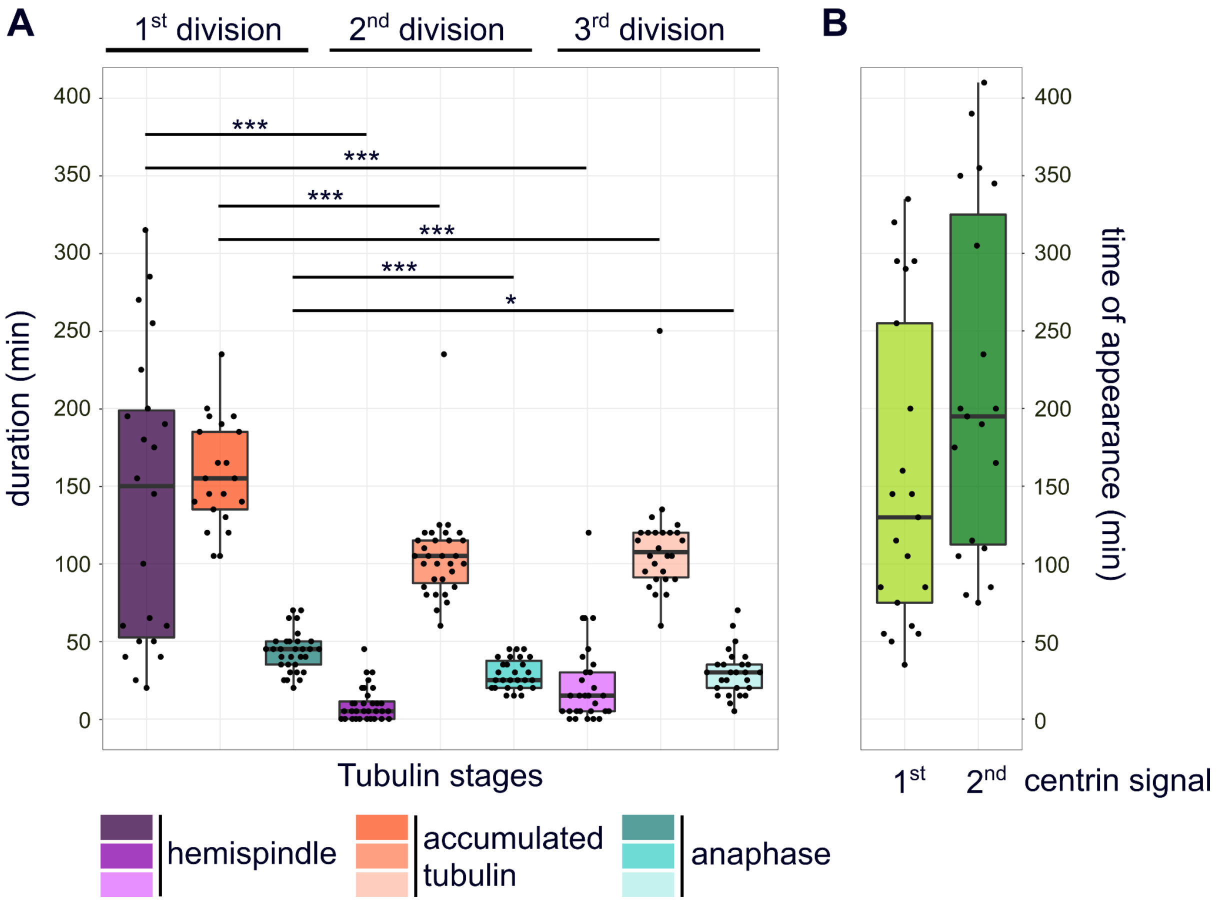


**Supplemental figure 1) Quantification of microtubule and centrin dynamics during the first three rounds of nuclear division.** A) Durations of subsequent microtubule organization phases in dividing blood stage parasites, i.e. hemispindle, accumulation, and anaphase, in 36 movies as shown in Fig. 1A. Since most movies (32/36) were already started during hemispindle phase, we quantified the minimal mean length of hemispindle stage for the first division. To test for significant differences we employed Welch Anova - Games-Howell test. B) Duration from start of movie until appearance of clear first and second PfCentrin1-GFP signals could be determined for 18 movies.

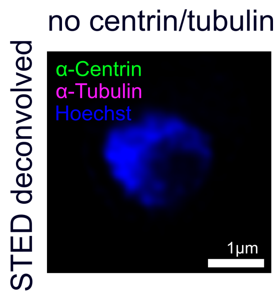


**Supplemental figure 2) Blood stage parasites show no tubulin or centrin staining in stages preceding schizogony.** Dual-color STED nanoscopy images of pre-schizogony trophozoite-stage parasite immunolabeled with anti-centrin (green), anti-tubulin (magenta) antibodies and stained with Hoechst (blue). Scale bar, 1 μm.

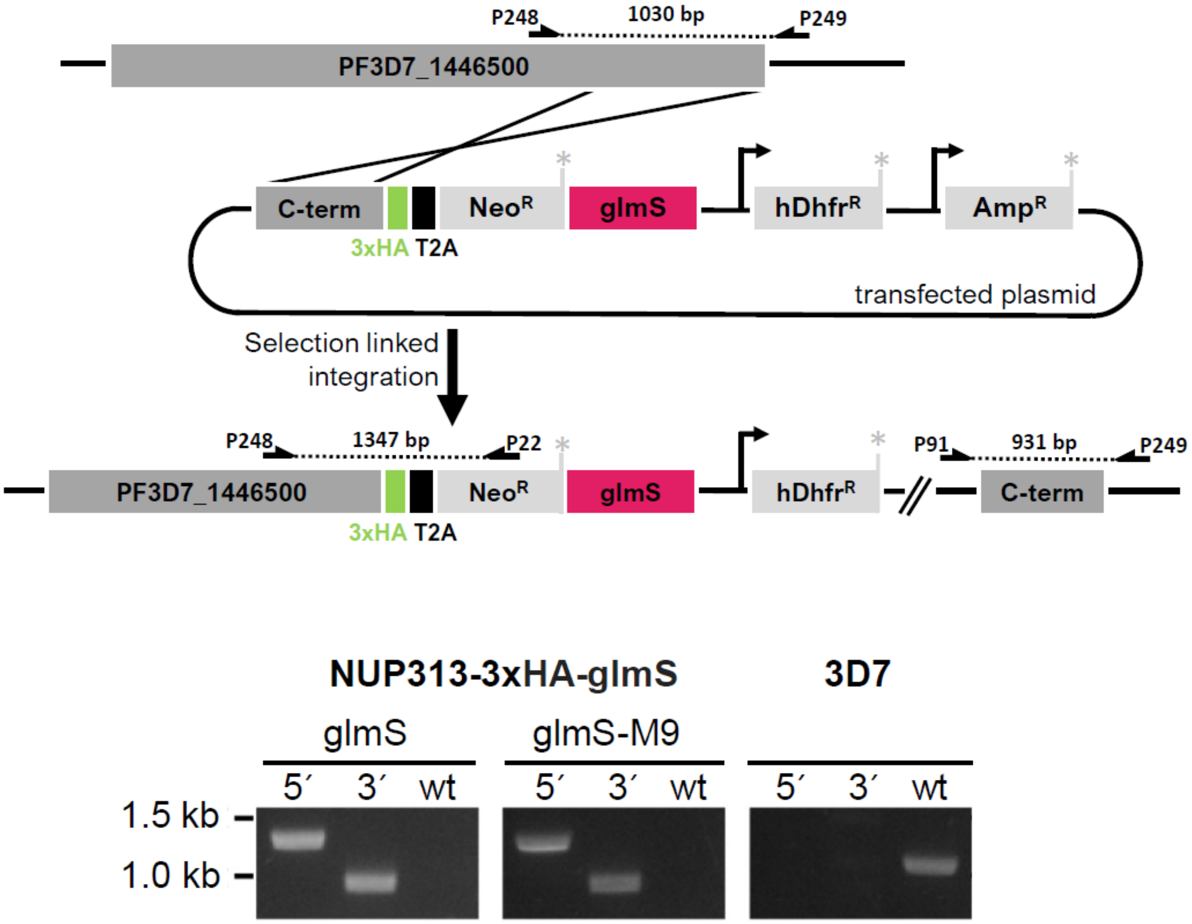


P22_KanR_seq_rev: CGCTTCAGTGACAACGTCGAGCACAGC
P91_3'_int_test_for: CACACAGGAAACAGCTATGACC
P248_NUP313_5' int_test_for: AGATCTGATTCCATTTCTGG
P249_NUP313_3' int_test_rev: GAGATAAGTAAGGATATACTTTTGC

**Supplemental figure 3) Tagging strategy of endogenous Nup313 with 3xHA and glms tag using selection linked integration.** The 3´-end of the open reading frame of the PF3D7_1446500 gene was cloned in pSLI-TGD-HA-glms construct to allow recombination with endogenous locus inducing expression of Neomycin selection cassette. PCRs using indicated primers show successful 5´ and 3´ integration into the genome and the absence of wild type locus after secondary selection. Two versions of the glms tag were cloned but only the original glms version was used in this study.

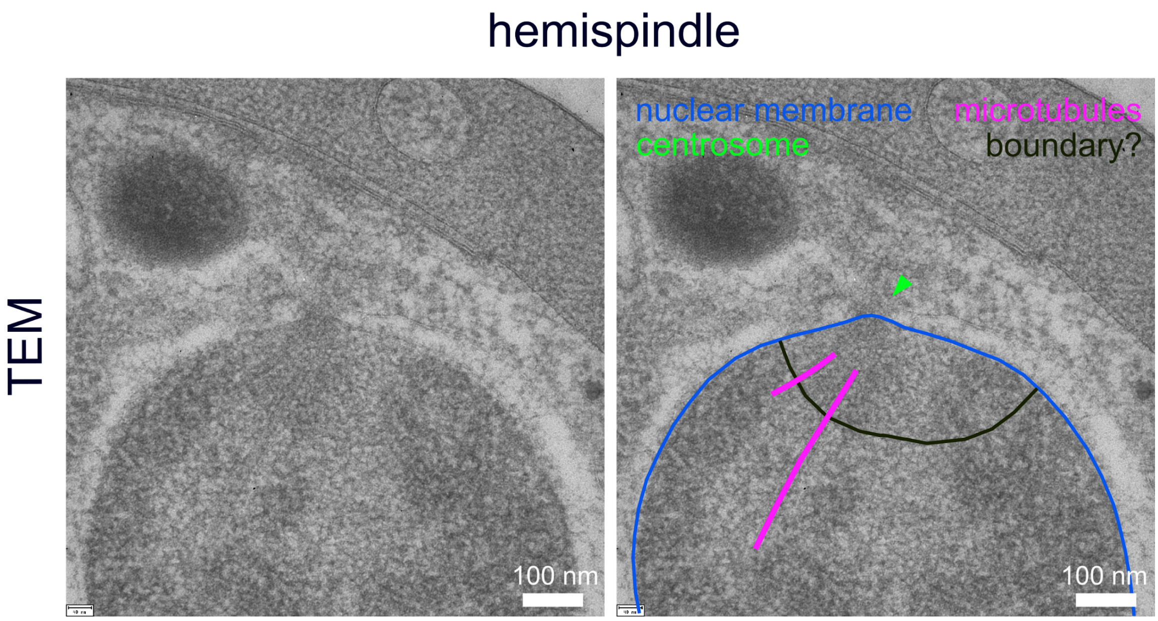


**Supplemental figure 4) Transmission electron microscopy reveals a microtubule-associated boundary region within the nucleus.** Transmission electron image of the nuclear region (annotated copy on the right) in a schizont shows no invagination of the nuclear membrane (blue) but suggests a boundary-like structure (black) delineating an intranuclear region from which microtubules (magenta) emanate. Arrow indicates likely positioning of the centriolar plaque (green). Scale bars are as indicated in image.

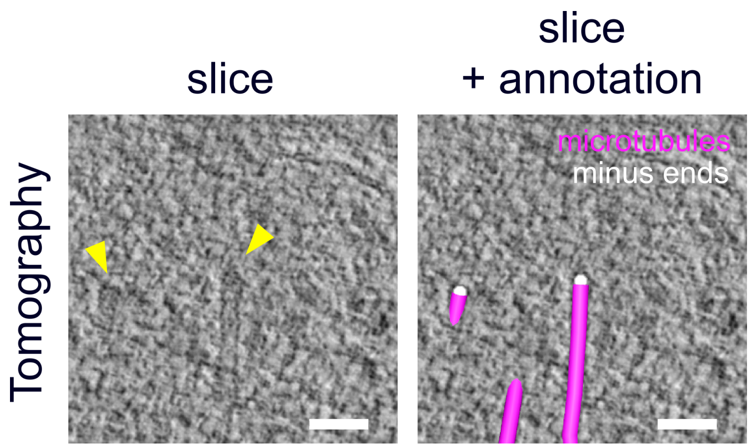


**Supplemental figure 5) Microtubule nucleating complex identified by minus end caps.** 3D-reconstructed electron tomography images of thick-sections (200nm) of high-pressure frozen and embedded schizonts showing the pole of a mitotic spindle. Arrows (yellow) indicate cap-like structures marking the minus ends of two microtubules of a tomogram slice on the left. Same image with manual annotations of microtubules (magenta) and minus ends (white) on the right. Scale bars, 50 nm.

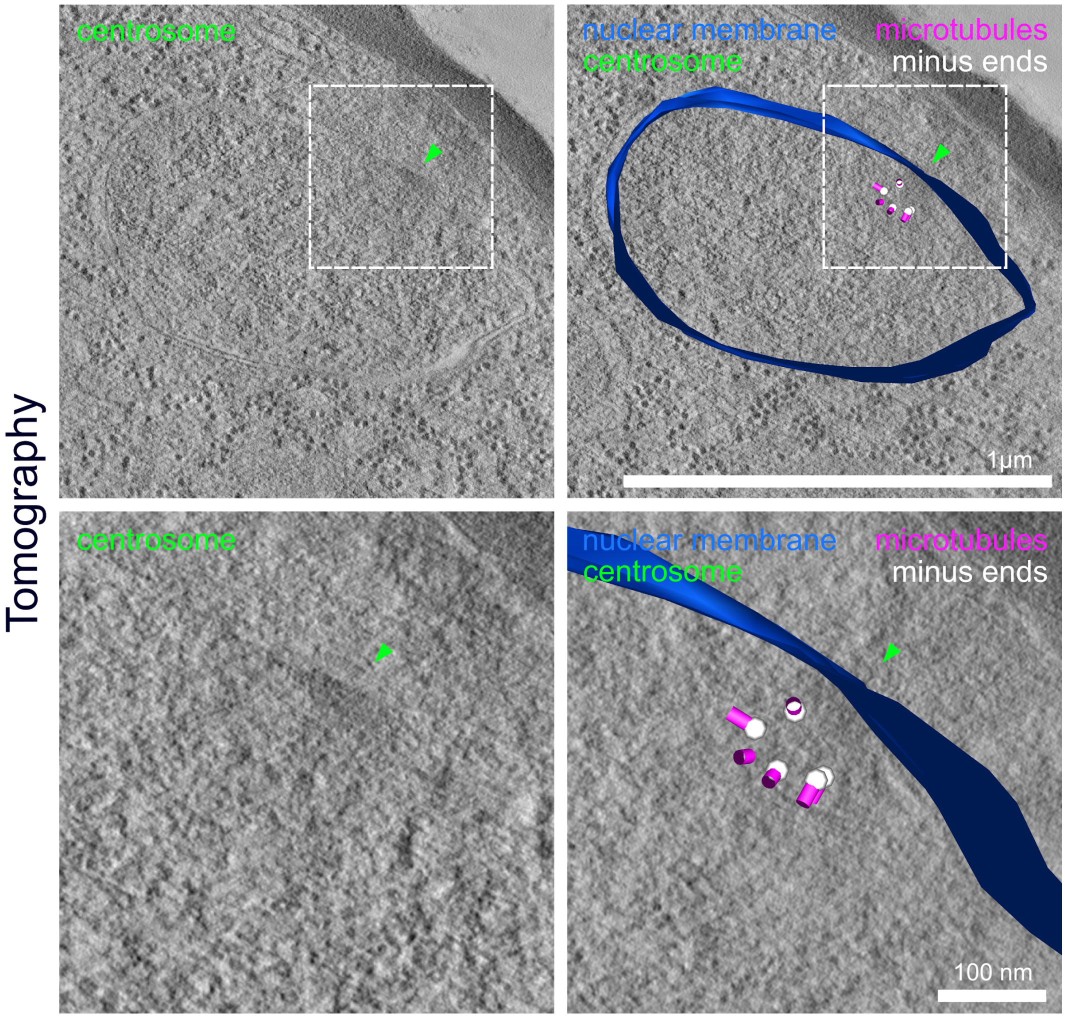


**Supplemental figure 6) Intranuclear region can be surmised in some electron tomographic sections.** 3D-reconstructed electron tomography images of thick-sections (200nm) of a high-pressure frozen and embedded schizont with zoom-ins. Arrows indicate likely positioning of the centriolar plaque (green). Microtubules (magenta), minus ends (white), nuclear membrane (blue) were manually annotated. Scale bars are indicated in images.

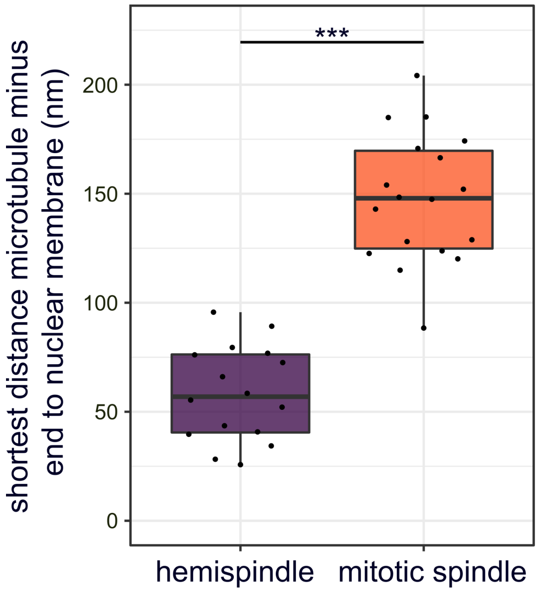


**Supplemental figure 7) Minus ends of intranuclear microtubules localize at a significant distance from the nuclear envelope.** Quantification of shortest distance between nuclear envelope and minus ends annotated in 3D-reconstructed electron tomography images of mitotic spindle (n=1, Fig. 3B) or hemispindles (n=4 Fig. S6, 3A). Unpaired t-test indicates significant difference.

#### Supplementary movies

Download under: https://heibox.uni-heidelberg.de/d/dcbfdb88b6e7492fbfaa/ . Please note that movies have been compressed to reduce file size (especially tomograms).

**Supplementary Movie 1) Live cell microscopy of microtubule and centriolar plaque dynamics.** Deconvolved confocal time-lapse microscopy movie of first spindle formation and elongation in a schizont ectopically expressing PfCentrin1-GFP (green) and labeled with SPY555-Tubulin (magenta). Maximum intensity projections are shown. Time interval is 5 min. (422 MB).

**Supplementary Movie 2) Live cell microscopy of microtubule and centriolar plaque dynamics.** As in supplementary movie 2 but with a late appearance of PfCentrin1-GFP signal. (22 MB).

**Supplementary Movie 3) Three dimensional organization of microtubules and centiolar plaques in dividing nuclei using ultrastructure expansion microscopy**. A) Slice-by-slice animation of a deconvolved confocal z-stack (17x300nm) acquired of a hemispindle containing nucleus in a Nup313-HA expressing schizont parasite labeled with anti-centrin (green), anti-tubulin (magenta), anti-HA (yellow) antibodies and stained with Hoechst (blue) after isotropic expansion by a factor of 4.5. (1 MB) B) 3D-rendering of the same nucleus using a Maximum Intensity Projection (MIP) mode in Imaris was used. Scale bars: 1 µm. (11 MB)

**Supplementary Movie 4) Three dimensional organization of microtubules and centriolar plaques in dividing nuclei using ultrastructure expansion microscopy.** As in supplementary movie 2 but showing nucleus with mitotic spindle. A) (2 MB). B) (10 MB).

**Supplementary Movie 5) Positioning of microtubule nucleation sites in hemispindle stage nucleus.** Slicing through 3D electron tomograms of thick-sections (200nm) of schizont nucleus in hemispindle stage. Corresponding surface rendering of microtubules (magenta), nuclear membrane (blue), microtubule minus ends (white), are animated subsequently. (338 MB).

**Supplementary Movie 6) Positioning of microtubule nucleation sites in mitotic spindle stage nucleus.** As in supplementary movie 4 for mitotic spindle stage. (357 MB).
